# DRP1-mediated mitochondrial fission integrates growth hormone signaling with metabolic and stress adaptation in triple-negative breast cancer

**DOI:** 10.64898/2026.03.08.710368

**Authors:** Juliana Moreira Mendonça-Gomes, Mariana Tominaga Pereira, Luísa Menezes-Silva, Luís Eduardo Duarte Gonçalves, Mariana Abrantes do Amaral, Lais Cavalieri Paredes, Camila Morales Fénero, Barbara Nunes Padovani, Mario Costa Cruz, Niels Olsen Saraiva Câmara

## Abstract

Triple-negative breast cancer (TNBC) relies on metabolic plasticity to sustain growth under diverse microenvironmental conditions. Although growth hormone (GH) signaling has been linked to breast cancer progression, its mechanistic integration with mitochondrial dynamics and metabolic reprogramming remains unclear. Here, we show that GH promotes TNBC progression accompanied by DRP1-associated mitochondrial remodeling, as indicated by sensitivity to Mdivi-1. The MDA-MB-231 line enabled integrated assessment across 2D, 3D, hypoxia, and in vivo xenografts using consistent workflows and readouts. GH increased proliferation and mitochondrial mass without increasing OCR under protein normalization. Instead, GH selectively enhanced glycolytic flux and metabolic flexibility. Inhibition of DRP1 uncoupled GH-induced glycolysis from proliferation, demonstrating that mitochondrial fission is required to link metabolic reprogramming to cell-cycle progression. DRP1 inhibition with Mdivi-1 was associated with altered TP53 and HIF1A expression and extended GH activity to the regulation of a pro-inflammatory tumor microenvironment marked by *cxcr4b*, *il8*, and *il12.* Consistent with these findings, analysis of human TNBC transcriptomes revealed conserved enrichment of mitochondrial, metabolic, and inflammatory pathways. Together, these results support the GH–DRP1 axis as a candidate regulator of mitochondrial dynamics, metabolic plasticity, tumor progression and tumor microenvironment interactions in TNBC.

## Introduction

Growth hormone (GH) is a central regulator of systemic growth and metabolism, exerting its effects through the GH receptor (GHR) and downstream activation of the JAK2–STAT5, PI3K–AKT, and MAPK pathways ^1,2^. While GH signaling has been extensively studied in the context of development and endocrine regulation, accumulating evidence suggests that it may also contribute to tumor initiation and progression ^3^. Elevated GH and insulin-like growth factor 1 (IGF-1) levels have been associated with increased risk of breast, prostate, and colorectal cancers, supporting a role for GH/IGF signaling in oncogenesis ^3,4^. However, the cellular and molecular mechanisms by which GH directly influences tumor cell metabolism and the tumor microenvironment remain poorly understood.

A hallmark of cancer cells is their ability to reprogram energy metabolism to support uncontrolled growth and survival under stress ^5,6^. One major adaptation is the shift toward glycolysis, even under normoxic conditions, known as the Warburg effect ^7^. This metabolic rewiring provides both rapid ATP production and biosynthetic precursors required for proliferation. Recent studies have revealed that mitochondrial dynamics, particularly the balance between fusion and fission, play a crucial role in regulating this metabolic flexibility ^8^. Excessive mitochondrial fission, mediated by the GTPase dynamin-related protein 1 (DRP1), has been linked to increased glycolysis, stress resistance, and enhanced tumorigenic potential ^9–11^. Conversely, mitochondrial fusion proteins such as mitofusin 2 (MFN2) are often downregulated in cancer, further skewing the balance toward fission and metabolic plasticity ^12,13^.

Stress-responsive pathways, including p53 and hypoxia-inducible factor 1α (HIF1α), are intimately connected to both mitochondrial remodeling and metabolic reprogramming ^14,15^. p53, beyond its canonical tumor suppressor functions, modulates glycolysis and oxidative metabolism depending on cellular context, while HIF1α orchestrates the transcriptional program for hypoxia adaptation, including glycolytic gene expression ^10,14,15^. These pathways provide tumors with a survival advantage in nutrient- and oxygen-deprived environments, such as the hypoxic tumor microenvironment ^14,16^.

Although GH has been implicated in the regulation of systemic metabolism, its direct effects on mitochondrial dynamics, tumor bioenergetics, and stress signaling remain largely unexplored. Given the established links between GH signaling, metabolic regulation, and cancer risk, we hypothesized that GH may act as a potent modulator of tumor cell metabolism through mitochondrial remodeling.

In this study, we investigated the effects of recombinant human GH (rhGH) on mitochondrial morphology, glycolytic and respiratory metabolism, and stress signaling in triple-negative breast cancer (TNBC) MDA-MB-231 cells. We combined 2D cultures, 3D spheroids, and zebrafish xenografts to evaluate the impact of rhGH in physiologically relevant models. Here, we show that growth hormone (GH) promotes triple-negative breast cancer progression by engaging DRP1-dependent mitochondrial remodeling that rewires glycolytic metabolism and stress-response pathways without increasing OCR under protein normalization. We further demonstrate that mitochondrial fission is required to couple GH-induced glycolysis to effective proliferation and to shape a pro-inflammatory tumor microenvironment characterized by cxcr4b, il8, and il12 induction. These findings uncover a previously unrecognized role of GH in tumor metabolic plasticity and identify mitochondrial fission as a critical mediator of GH-driven cancer progression.

## Materials and Methods

### 1. Cell Lines and Culture Conditions

For this study, the breast cancer cell line MDA-MB-231 mKate2 (triple-negative) was used. The cells were kindly donated by Professor Sonia Jancar Negro (USP, São Paulo, SP, Brazil) and maintained under standard culture conditions. Cells were grown in Dulbecco’s Modified Eagle Medium (DMEM; Thermo Fisher Scientific, Waltham, MA, USA) supplemented with 10% fetal bovine serum (FBS; Corning, Corning, NY, USA) and 1% penicillin-streptomycin (Gibco, Grand Island, NY, USA). Cultures were incubated at 37 °C in a humidified atmosphere containing 5% CO□. Cells were subcultured upon reaching approximately 70–80% confluence. For passaging, cultures were washed with phosphate-buffered saline (PBS; Thermo Fisher Scientific) to remove residual medium and detached using 0.25% trypsin-EDTA solution (Gibco™). After detachment, cells were resuspended in fresh culture medium, counted using a hemocytometer, and seeded at appropriate densities for each experimental condition.

### 2. 3D Spheroid Culture and Magnetic Manipulation

MDA-MB-231 mKate2 cells cultivated under standard 2D conditions, as described above, were passaged every 2–3 days to maintain optimal growth. For spheroid formation, labeled cells (1 × 10□ cells/well) were seeded in 96-well Ultra-Low Attachment Plates (CellStar®, Greiner Bio-One) in 150 µL of complete medium. These plates promote the self-assembly of cells into spheroids due to their low-attachment surface. Cultures were incubated at 37 °C for 48 h to allow spheroid aggregation. To enable magnetic manipulation, a permanent magnet array (Greiner Bio-One) was positioned beneath the culture plate. A magnetic field (0.1–1.0 Tesla) was applied for 30 min to guide and align spheroids within the wells. The magnetic field was reapplied every 48 h to reposition spheroids and maintain uniform organization. Cultures were kept in a humidified incubator at 37 °C with 5% CO□.

### 3. Growth Hormone Treatment (rhGH)

To investigate the effects of growth hormone on mitochondrial dynamics and metabolic reprogramming, MDA-MB-231 cells grown in both monolayer and 3D cultures were treated with recombinant human GH (rhGH; PeproTech, Rocky Hill, NJ, USA) at a final concentration of 100 ng/mL. The hormone was diluted in DMEM to the desired concentration, and treatments were applied for 48, 96, or 144 h, depending on the experimental setup. Control groups received DMEM without GH.

### 4. Inhibition of Mitochondrial Fission

To further investigate the role of mitochondrial fission in GH-induced metabolic reprogramming, both 2D and 3D cultures were treated with the mitochondrial fission inhibitor Mdivi-1 (Sigma-Aldrich, St. Louis, MO, USA) at a final concentration of 15 µM for 24 h following GH treatment. Control cells received an equivalent amount of dimethyl sulfoxide (DMSO) as a vehicle.

### 5. Hypoxia induction

Hypoxic conditions were established using a modular Hypoxia Incubator Chamber (StemCell Technologies). Cells were cultured under 1% O□, 5% CO□, and 94% N□ at 37 °C in a humidified atmosphere. For hypoxia exposure, culture plates were placed in the chamber, flushed with the gas mixture for 5 min at a flow rate of 20 L/min, and sealed according to the manufacturer’s instructions. Cells were maintained under hypoxia for 24h, depending on the experimental setup, while normoxic controls were cultured in standard incubators (21% O□, 5% CO□).

### 6. Mitochondrial Morphology Analysis

Mitochondrial morphology was assessed by fluorescence imaging. Cells were stained with MitoTracker Green (Thermo Fisher Scientific) for 30 min at 37 °C and washed with PBS. Images were acquired using a confocal laser scanning microscope Zeiss LSM-780-NLO (Carl Zeiss, Germany), and mitochondrial morphology was analyzed with Imaris software (Bitplane, Zurich, Switzerland). Mitochondrial fission was quantified by counting the number of fragmented mitochondria per cell.

### 7. Glucose and lactate quantification

Glucose and lactate levels were measured in 96-well flat-bottom microplates (Corning Inc., NY, USA), with separate plates designated for each metabolite. Samples were homogenized by vortexing and centrifuged at 1500 rpm for 5 min. Aliquots of 2.5 µL from each sample were transferred into the corresponding wells. Blank controls contained 2.5 µL of distilled water, and calibration was performed using 2.5 µL of standard solutions prepared according to the manufacturer’s instructions (Labtest, Brazil). Following sample and control preparation, 250 µL of the respective assay reagent was added to all wells, including blanks and standards. The glucose plate was incubated at 37 °C for 10 min, and the lactate plate for 5 min under the same conditions. Absorbance was recorded at 505 nm using a Synergy H1 microplate reader (BioTek Instruments, Winooski, VT, USA). Data acquisition and analysis were carried out with Gen5 software (BioTek Instruments).

### 8. Protein Expression (Western Blot)

Cells were lysed in RIPA buffer (Sigma-Aldrich) supplemented with protease and phosphatase inhibitor cocktail (Thermo Fisher Scientific). Protein concentration was determined using the bicinchoninic acid (BCA) assay (Sigma-Aldrich). Equal amounts of protein (30 µg) were mixed with 2× Laemmli sample buffer (Bio-Rad), heated at 95 °C for 10 min, and resolved on 12% gradient polyacrylamide gels (Bio-Rad). Proteins were transferred onto 0.2 µm nitrocellulose membranes (Bio-Rad) using the Trans-Blot Turbo system (Bio-Rad). Membranes were blocked with 5% non-fat milk in TBST for 1 h and incubated overnight at 4 °C with primary antibodies against target proteins, using housekeeping proteins as loading controls. After incubation with HRP-conjugated secondary antibodies, bands were visualized by enhanced chemiluminescence (ECL) with the ChemiDoc™ XRS imaging system (Bio-Rad) and quantified using Image Lab 3.0 software (Bio-Rad).

### 9. RNA Extraction and Quantitative Real-time PCR (qRT-PCR)

MDA-MB-231 cells cultured with or without recombinant GH were processed for RNA extraction using the ReliaPrep™ RNA Extraction Kit (Promega) according to the manufacturer’s instructions. RNA concentration and purity were assessed with a NanoDrop spectrophotometer (Thermo Fisher Scientific, Waltham, MA, USA). For cDNA synthesis, 2 µg of RNA was reverse-transcribed, and the resulting cDNA was diluted 1:10 in nuclease-free water prior to analysis. qRT-PCR reactions were performed with 4 µL of diluted cDNA, 0.5 µL of each primer (final concentration 125–500 nM), and 5 µL of PowerUp™ SYBR Green Master Mix (Thermo Fisher Scientific, USA) using a QuantStudio™ 12K Flex PCR System (Applied Biosystems). β-actin was used as the reference gene. Relative gene expression was calculated using the 2□ΔΔCT method. Primer sequences are listed in the Primer sequences are listed in Table 1. No amplification was observed in no-template controls, and all primers displayed a single melting curve peak with efficiencies >90%.

### 10. Metabolic Flux Analysis (OCR and ECAR)

To assess metabolic function in MDA-MB-231 cells in the presence or absence of recombinant GH, oxygen consumption rate (OCR) and extracellular acidification rate (ECAR) were measured using the XF96 Seahorse Extracellular Flux Analyzer (Agilent Technologies, USA), which evaluates mitochondrial respiration and glycolysis, respectively. Cells (2 × 10□/well) were seeded on poly-L-lysine–coated XF96 cell culture plates. The day prior to the assay, sensor cartridges were hydrated according to the manufacturer’s protocol. One hour after seeding at 37 °C, cells were washed twice with Seahorse XF minimal DMEM (Agilent) supplemented with 1 mM glutamine (Gibco), pH 7.4, and incubated for 1 h at 37 °C in a CO□-free incubator. For OCR assays, cells were maintained in Seahorse XF minimal DMEM supplemented with 25 mM glucose (Sigma), 1 mM pyruvate (Sigma), and 1 mM glutamine. After calibration, basal respiration was measured, followed by sequential injections of 1 µM oligomycin (ATP synthase inhibitor), 0.5 µM FCCP (uncoupler to assess maximal respiratory capacity), and 1 µM rotenone plus 1 µM antimycin A (complex I and III inhibitors, respectively) to determine non-mitochondrial respiration. For ECAR assays, the medium contained Seahorse XF minimal DMEM supplemented with 1 mM glutamine. Following calibration, sequential injections were performed: 10 mM glucose to initiate glycolysis, 1 µM oligomycin to increase glycolytic flux, and 50 mM 2-deoxy-D-glucose (2-DG) to inhibit glycolysis. All respiratory modulators were pre-titrated according to manufacturer recommendations. After each assay, protein content was quantified from the same wells for normalization. Data acquisition and analysis were performed using Wave software (Agilent Technologies, USA).

### 11. Zebrafish line and maintenance

#### a. Ethical Statement

All animal experiments were approved by the Ethics Committee for Animal Use (CEUA-ICB/USP; protocol no. 9710200720) and conducted in accordance with the Brazilian Guidelines for Care and Use of Animals for Scientific and Teaching Purposes (CONCEA) as well as international guidelines for animal experimentation.

#### b. Zebrafish Husbandry and Ethics

Adult zebrafish (*Danio rerio*) of the AB wild-type strain and the Tg(GH:GFP) transgenic line were maintained in a recirculating aquarium system at 28–28.5 °C under a 14 h light/10 h dark cycle at the Zebrafish Facility, Institute of Biosciences, University of São Paulo. Fish were housed at a density of 5 fish/L in aerated, dechlorinated water and fed twice daily with Zeigler Larval Diet (AP450, Zeigler Bros., USA) and supplemented with live brine shrimp (*Artemia nauplii*). For experiments, Tg(GH:GFP) fish ^17^ were outcrossed with AB wild-type animals to generate GH:GFP□ and GH:GFP□ sibling controls. Embryos were collected and maintained in E3 medium (5 mM NaCl, 0.17 mM KCl, 0.33 mM CaCl□, 0.33 mM MgSO□) supplemented with 0.05% methylene blue, and transferred to nursery tanks at 5 days post-fertilization (dpf). Larvae were fed live feed and gradually transitioned from AP50 to AP450 diets (Zeigler) according to developmental stage.

#### c. Zebrafish Xenograft Assay

MDA-MB-231 cells were xenografted into adult zebrafish (4–6 months old) via intraperitoneal injection, a well-established method for tumor engraftment in zebrafish models. Fish were anesthetized in system water containing 0.04% tricaine (MS-222; Sigma-Aldrich), and a small incision was made near the dorsal region using a sterile scalpel. A Hamilton syringe fitted with a 30-gauge needle (Hamilton Company) was used to inject 2–3 µL of cell suspension (2 × 10□ cells) into the perivitelline cavity. Following injection, zebrafish were transferred to recovery tanks with fresh system water for 24 h before being returned to the main aquarium. Animals were monitored daily for signs of distress, infection, or abnormal behavior; those exhibiting adverse effects were excluded from the study.

#### d. Tumor Monitoring

Tumor growth was monitored for up to 2 weeks post-injection. Tumor formation was assessed every 3–4 days by visual inspection under a fluorescence or brightfield microscope (ZEISS Axio Zoom.V16). Fluorescence imaging was performed in xenografts generated with GFP- or RFP-labeled cells to facilitate visualization. For non-labeled tumors, growth was evaluated based on external mass protrusion or morphological changes in the zebrafish.

#### e. Fluorescence Imaging of Tumors

For monitoring fluorescent tumors, zebrafish were imaged using either a fluorescence stereomicroscope (ZEISS Axio Zoom.V16) at defined time points. Images were acquired to track tumor growth, and fluorescence intensity was quantified with ImageJ software (NIH) to estimate tumor size over time.

### 12. Statistical Analysis

All experiments were performed in triplicate, and results are expressed as mean ± standard deviation (SD). Statistical analyses were conducted using GraphPad Prism 10.6.1 (GraphPad Software, San Diego, CA, USA). Group differences were evaluated by one-way ANOVA followed by Sidak’s post hoc test or by Student’s *t*-test, as appropriate. A *p*-value < 0.05 was considered statistically significant.

## Results

### 1. rhGH Promotes Proliferation with Concurrent Mitochondrial Remodeling and Stress Activation

To evaluate the effect of recombinant human growth hormone (rhGH) on breast cancer cell proliferation and metabolism, MDA-MB-231 cells were cultured for six days in the presence or absence of rhGH (Fig. 1A). GH-treated cells showed a significant increase in total cell number compared to controls (Fig. 1B), as well as higher total protein content (Fig. 1C), indicating enhanced proliferative activity. MitoTracker Green staining demonstrated that GH treatment led to a marked increase in mitochondrial mass, as evidenced by quantitative image analysis of fluorescence intensity and confirmed by flow cytometry through increased mean fluorescence intensity (MFI) (Fig. 1D–E). Western blot analysis revealed higher levels of the mitochondrial fission protein DRPI and its phosphorylated form (pDRP1) in GH-treated cells, while MFN2 protein levels remained stable (Fig. 1F). At the transcriptional level, qPCR analyses demonstrated lower expression of *DNM1L* after six days of GH treatment (Fig. 1G). However, at earlier points (Fig. S1A), *DNM1L* expression was transiently upregulated, particularly at day 4, suggesting a temporal regulation and a possible early trigger preceding the increase in DRP1 protein levels observed at day 6. Regarding *MFN2*, its mRNA expression was reduced at day 6 (Fig. 1G) and remained unchanged during the first two points (Fig. S1A). Similarly, *TP53* and *HIF1A* transcripts, which were downregulated after two days of GH exposure, remained stable at day 4 (*HIF1A)* (Fig. S1B) and were increased at six days for both genes (Fig. 1G), indicating an early adaptive response to GH-induced metabolic stress. Together, these results indicate that GH promotes MDA-MB-231 cell proliferation and increases mitochondrial mass, accompanied by dynamic, time-dependent transcriptional and post-transcriptional changes in genes involved in mitochondrial remodeling and cellular stress adaptation.

**Figure 1.**
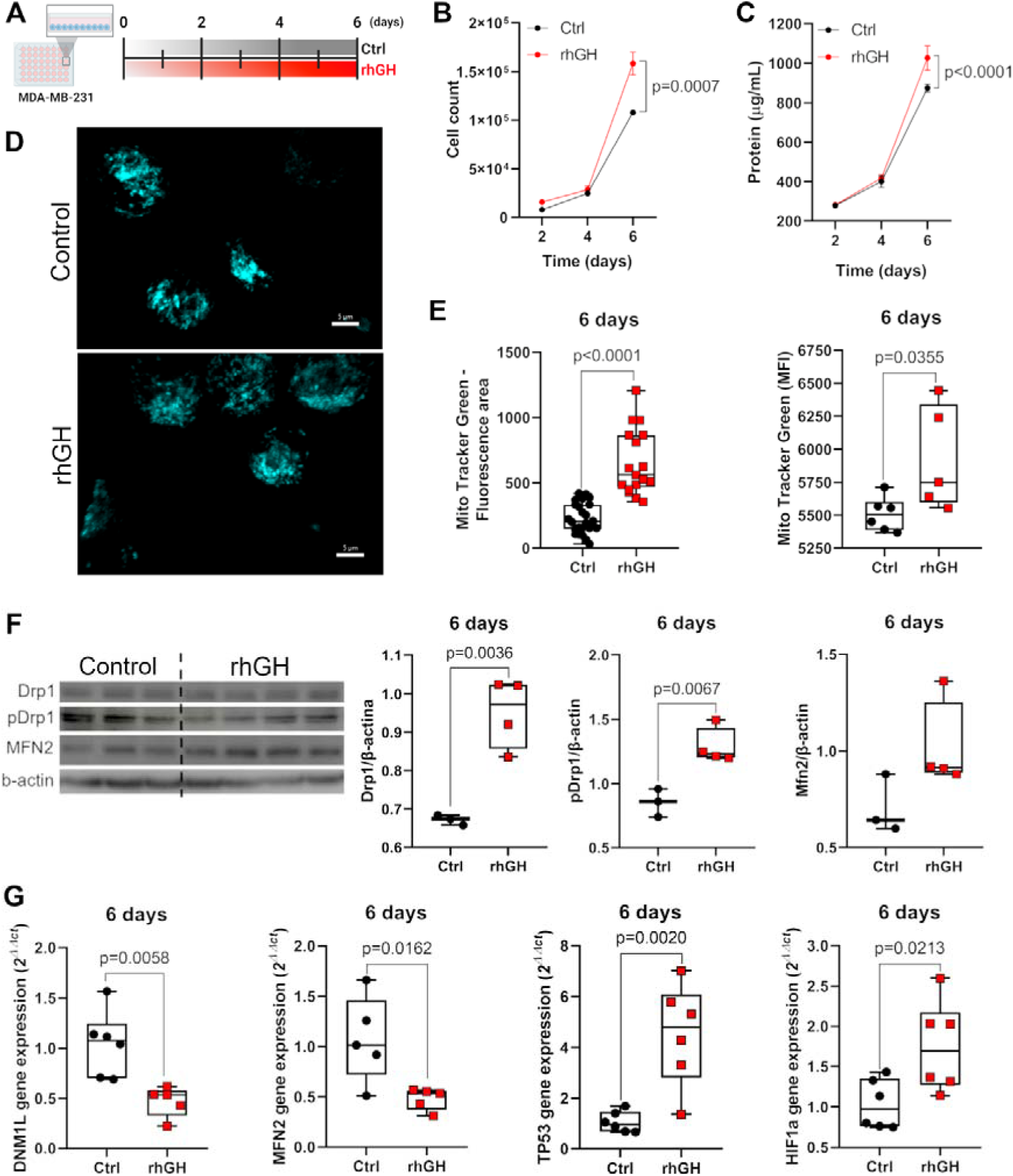
Growth hormone enhances proliferation and induces mitochondrial remodeling and stress-response signaling in MDA-MB-231 breast cancer cells. (A) Experimental timeline showing MDA-MB-231 cells cultured for up to 6 days under control conditions or treated with recombinant human growth hormone (rhGH). (B) Cell counts over time and (C) Total protein content (µg/mL). (D) Representative confocal images of mitochondrial networks stained with MitoTracker Green (cyan) in control and rhGH-treated cells after 6 days. Scale bars, 5 µm. (E) Quantification of MitoTracker Green fluorescence expressed as total fluorescence per area (left) and mean fluorescence intensity (MFI; right). (F) Representative immunoblots and densitometric analyses of DRP1, phosphorylated DRP1 (pDRP1), and MFN2 normalized to β-actin after 6 days of treatment. (G) Relative mRNA expression levels of DNM1L, MFN2, TP53, and HIF1A after 6 days of culture. Data are presented as mean ± SEM. Statistical significance was determined using an unpaired Student’s *t*-test or two-way ANOVA followed by Sidak’s post hoc test.

### 2. Growth hormone enhances glycolytic metabolism without altering mitochondrial respiration in MDA-MB-231 cells

To investigate whether GH-induced proliferation and increased mitochondrial mass were associated with metabolic changes, we performed Seahorse extracellular flux analyses in MDA-MB-231 cells cultured with or without recombinant human GH (rhGH) for 2, 4, and 6 days. At day 6, GH-treated cells displayed significantly higher extracellular acidification rates (ECAR), indicating enhanced glycolytic activity (Fig. 2A, S2A). Both glycolysis and glycolytic capacity were increased at 4 and 6 days, respectively (Fig. S2B and 2B), whereas the glycolytic reserve was consistently lower at 4 days and higher at 6 days (Fig. S2B and 2B). Both conditions exhibited similar glucose uptake rates throughout the time course (Fig. 2C), but GH-treated cells showed a marked accumulation of lactate at day 6 (Fig. 2C). Conversely, oxygen consumption rate (OCR) profiles demonstrated that GH treatment did not significantly alter OCR when normalized to total protein under the assay conditions (Fig. 2A, S2C). Basal respiration, ATP production, maximal respiration, proton leak, and spare respiratory capacity remained unchanged throughout the exposure period (Fig. S2D). Together, these data show increased extracellular acidification and lactate accumulation with unchanged OCR in intact-cell assays, supporting a shift toward glycolytic engagement under the conditions tested

**Figure 2.**
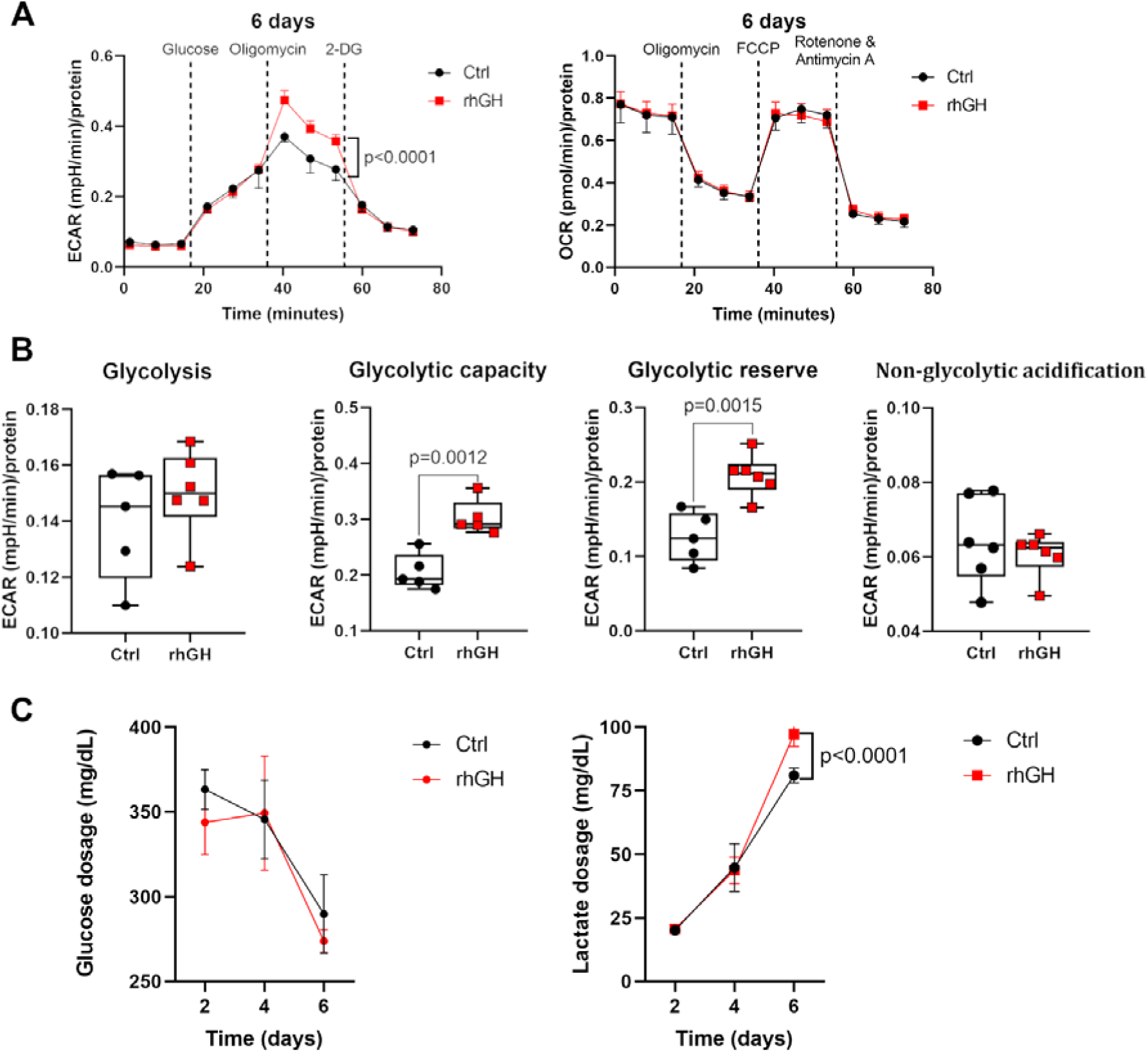
Growth hormone enhances glycolytic capacity and lactate production in breast cancer cells. (A) Representative extracellular flux analysis traces showing extracellular acidification rate (ECAR, left) and oxygen consumption rate (OCR, right) in control (Ctrl) and recombinant human growth hormone–treated (rhGH) MDA-MB-231 cells after 6 days of culture. ECAR was measured following sequential injections of glucose, oligomycin, and 2-deoxy-D-glucose (2-DG), while OCR was assessed after sequential injections of oligomycin, FCCP, and rotenone/antimycin A, as indicated. (B) Quantification of glycolytic parameters derived from ECAR measurements, including basal glycolysis, glycolytic capacity, glycolytic reserve, and non-glycolytic acidification. (C) Glucose consumption (left) and lactate production (right) measured in culture media over time (2, 4, and 6 days) in Ctrl and rhGH-treated cells. Data are presented as mean ± SEM with individual data points representing independent biological replicates. ECAR and OCR values were normalized to total protein content. Statistical significance was determined using a two-way ANOVA followed by Sidak’s post hoc test, and an unpaired Student’s *t*-test; *p* values are indicated in the graphs.

### 3. Inhibition of Mitochondrial Fission Modulates GH-Associated Mitochondrial and Glycolytic Responses

To determine whether mitochondrial fission contributes to the metabolic effects induced by growth hormone (GH), MDA-MB-231 cells were cultured for six days in the presence or absence of recombinant human growth hormone (rhGH). On day 5 of culture, both conditions received the DRP1 inhibitor Mdivi-1 (15 µM) for the final 24 hours, while GH exposure was maintained through day 6 (Fig. 3A). Confocal microscopy revealed that GH increased mitochondrial sphericity, consistent with a less elongated and more fragmented mitochondrial network, whereas Mdivi-1 markedly reduced mitochondrial sphericity even in the presence of GH, indicating a shift toward mitochondrial fusion and elongation (Fig. 3B and D). Mdivi-1 did not enhance proliferation when combined with GH, indicating that short-term DRP1 inhibition does not synergize with GH-mediated proliferative effects (Fig. 3C). Seahorse extracellular flux analysis showed that Mdivi-1 treatment modulated the extracellular acidification rate (ECAR), reversing the GH-associated changes in glycolytic capacity and glycolytic reserve while increasing basal glycolysis (Fig. 3E and G). Inhibition of DRP1 resulted in increased basal glycolytic activity both in the presence and absence of GH. Under GH exposure, DRP1 inhibition was associated with the maintenance of glycolytic capacity and glycolytic reserve at basal levels. Non-glycolytic acidification was not affected by either treatment (Fig. 3G). Conversely, oxygen consumption rate (OCR) parameters, including basal respiration, proton leak, ATP production, maximal respiration, spare respiratory capacity, and non-mitochondrial oxygen consumption remained unchanged by either GH or Mdivi-1 treatment (Fig. S3A-F). Together, these findings indicate that GH promotes mitochondrial fragmentation and enhances glycolytic metabolism in MDA-MB-231 cells through a DRP1-dependent mechanism. Short-term inhibition of mitochondrial fission by Mdivi-1 during sustained GH exposure altered GH-associated mitochondrial and metabolic responses and was associated with enhanced glycolysis, supporting a relationship between mitochondrial dynamics and metabolic regulation.

**Figure 3.**
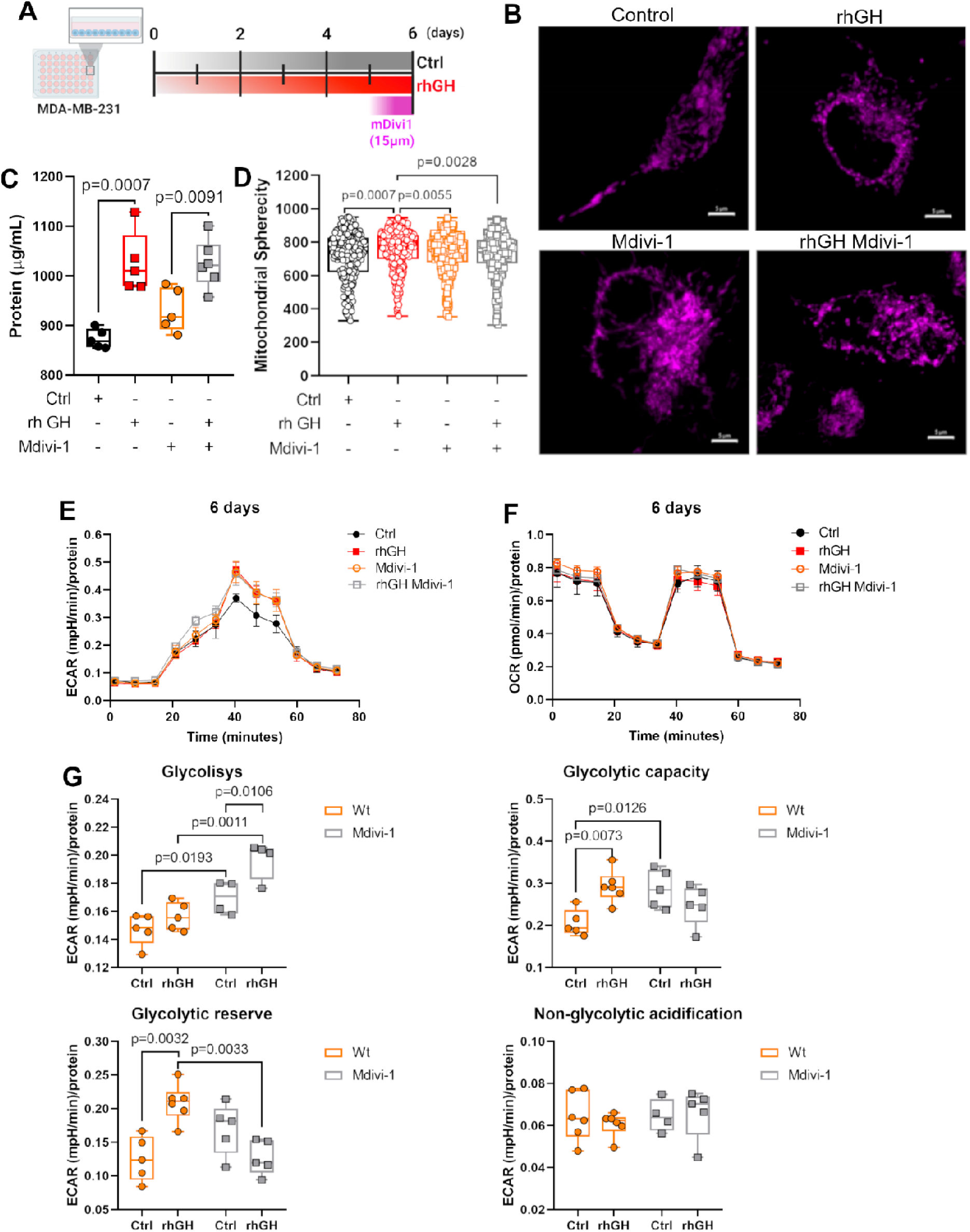
Growth hormone–driven mitochondrial remodeling and metabolic reprogramming are dependent on DRP1 activity in MDA-MB-231 cells. (A) Experimental timeline showing MDA-MB-231 cells cultured for 6 days under control conditions or treated with recombinant human growth hormone (rhGH), with the DRP1 inhibitor Mdivi-1 (15 µM) added as indicated. (B) Representative confocal images of mitochondrial networks (magenta) in control, rhGH-treated, Mdivi-1–treated, and combined rhGH + Mdivi-1 conditions. Scale bars, 10 µm. (C) Total protein content (µg/mL) after 6 days of treatment, as an indicator of cell proliferation following GH and Mdivi-1 treatments. (D) Quantification of mitochondrial sphericity. (E–F) Seahorse extracellular flux analysis after 6 days of treatment showing extracellular acidification rate (ECAR; E) and oxygen consumption rate (OCR; F) over time in response to sequential metabolic perturbations. (G) Quantification of glycolytic parameters derived from ECAR measurements, including glycolysis, glycolytic capacity, glycolytic reserve, and non-glycolytic acidification. Data are shown as individual values with mean ± SEM. Statistical significance was determined using a two-way ANOVA followed by Sidak’s post hoc test, with exact *p* values indicated.

### 4. Mdivi-1 modulates DRP1 activation and mitochondrial dynamics–related gene expression in GH-treated MDA-MB-231 cells

To further explore the molecular mechanisms underlying the metabolic effects of growth hormone (GH) and the role of mitochondrial fission, we examined the impact of DRP1 inhibition by Mdivi-1 on protein and gene expression in MDA-MB-231 cells. Cells were cultured as described previously, in the presence or absence of recombinant human growth hormone (rhGH), and treated with the DRP1 inhibitor Mdivi-1 (15 µM) for 24 hours following five days of GH exposure, with GH stimulation maintained throughout the experiment, for a total duration of six days (Fig. 4A). The increase in phosphorylated DRP1 (pDRP1) observed after six days of GH exposure was reversed by Mdivi-1 treatment, consistent with its role as an inhibitor of mitochondrial fission. Regarding MFN2, no significant changes were observed following Mdivi-1 treatment. Acetyl-p53 protein expression was reduced following DRP1 inhibition and was further decreased in the GH-treated group, suggesting a combined effect of GH and Mdivi-1 on the downregulation of acetyl-p53 levels. In contrast, HIF-1α protein levels were significantly reduced by Mdivi-1 in both control and GH-treated conditions to a similar extent, indicating that GH did not further modulate this response (Fig. 4B–C). At the transcriptional level, GH downregulated *DNM1L* and *MFN2* expression, and these reduced levels were maintained after Mdivi-1 treatment. In contrast, *TP53* and *HIF1A* transcripts increased by GH and were reduced in cultures treated with Mdivi-1, remaining lower when Mdivi-1 was combined with GH (Fig. 4D). The same experiment was also performed at an earlier point. Cells were cultured in the presence or absence of GH and treated with the DRP1 inhibitor Mdivi-1 (15 µM) for 24 hours following three days of GH exposure (4-day condition; Fig. S4A). Total protein content remained unchanged between groups, suggesting no increase in cell proliferation (Fig. S4B). At the early point, Mdivi-1 abolished GH-induced changes in DRP1 phosphorylation (pDRP1) in GH-treated cells. MFN2 protein levels remained unchanged among groups. Notably, acetyl-p53 expression was significantly increased by Mdivi-1 in GH-treated cells, while HIF1α levels were not affected (Fig. 4C–D). Together, these findings indicate that GH enhances DRP1 activation and induces transcriptional changes in mitochondrial dynamics and stress-related genes. Short-term inhibition of mitochondrial fission by Mdivi-1 reduces DRP1 phosphorylation and reverses GH-induced upregulation of *TP53* and *HIF1A*, highlighting the interplay between mitochondrial dynamics and stress-adaptive gene regulation.

**Figure 4.**
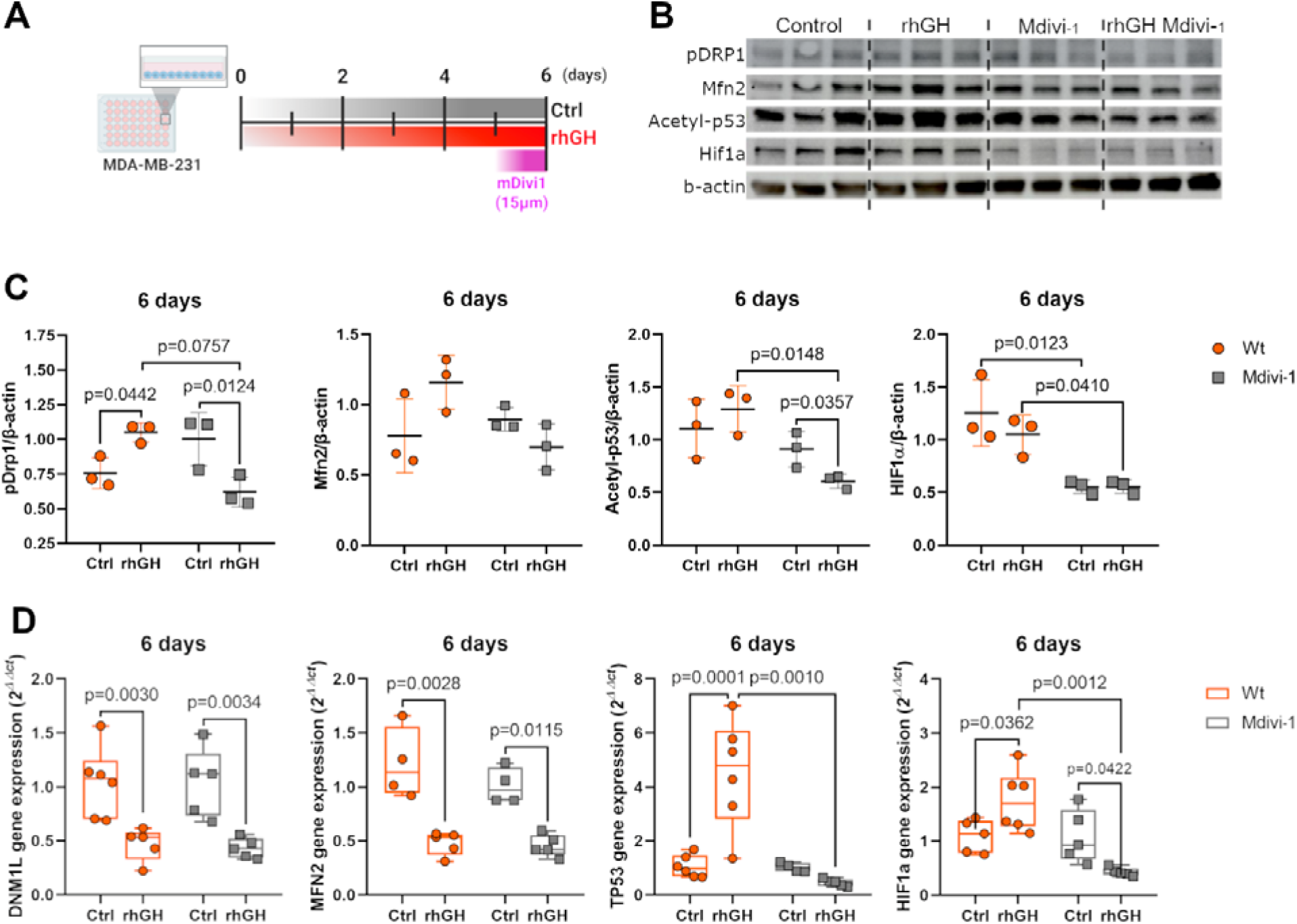
DRP1 inhibition modulates GH-associated mitochondrial signaling. (A) Experimental timeline illustrating MDA-MB-231 cell culture and treatment conditions. Cells were cultured for six days in the presence or absence of recombinant human growth hormone (rhGH). The DRP1 inhibitor Mdivi-1 (15 µM) was added during the final 24 hours while GH exposure was maintained throughout the experiment. (B) Representative immunoblots showing phosphorylated DRP1 (pDRP1), MFN2, acetyl-p53, and HIF-1α protein levels under the indicated conditions. β-actin was used as a loading control. (C) Quantification of protein levels normalized to β-actin after six days of treatment. (D) Relative mRNA expression levels of *DNM1L*, *MFN2*, *TP53*, and *HIF1A* after six days of treatment, assessed by qPCR and normalized to housekeeping genes. Statistical significance was determined using two-way ANOVA followed by Sidak’s post hoc test. Exact *p* values are indicated.

### 5. Growth hormone enhances proliferation and reshapes metabolic and stress-response pathways in MDA-MB-231 cells under hypoxia

To determine whether growth hormone (GH) influences cell metabolism and stress-response pathways under low-oxygen conditions, MDA-MB-231 cells were cultured for 1 day in 1% O□ in the presence or absence of recombinant human GH (rhGH) (Fig. 5A). Under hypoxic conditions, GH-treated cells exhibited increased cell number compared with hypoxic controls, as determined by total cell counts. Comparison of control samples under hypoxic versus normoxic conditions revealed a reduction in cell number under hypoxia, consistent with increased cell death in low-oxygen environments (Fig. 5B). This observation recapitulates features of solid tumors, which commonly exhibit a hypoxic core characterized by necrotic cells surrounded by viable tumor cells. At the transcriptional level, hypoxic conditions induced the overexpression of *DNM1L*, *TP53*, and *HIF1A*. However, when growth hormone (GH) was introduced in the hypoxic context, only *TP53* remained strongly upregulated, whereas the other genes were downregulated. These findings further highlight the central role of *TP53* in mediating GH responses under an additional stress condition, the hypoxia (Fig. 5C). Analysis of extracellular metabolites revealed that GH exposure increased glucose consumption per cell under hypoxic conditions. Nevertheless, overall glucose consumption remained lower in hypoxia compared with normoxia, with GH acting to potentiate glucose uptake rather than fully restore it. In contrast, lactate production was elevated under hypoxic conditions; however, GH treatment reduced lactate levels to near-basal values (Fig. 5D), suggesting a GH-mediated metabolic reprogramming under hypoxic stress. Together, these results indicate that even after a 1-day exposure to hypoxia, GH increases net cell number and glucose consumption, downregulates DNM1L and HIF1A, and induces *TP53* expression, suggesting that GH reshapes metabolic and stress-adaptive responses of breast cancer cells under oxygen restriction.

**Figure 5.**
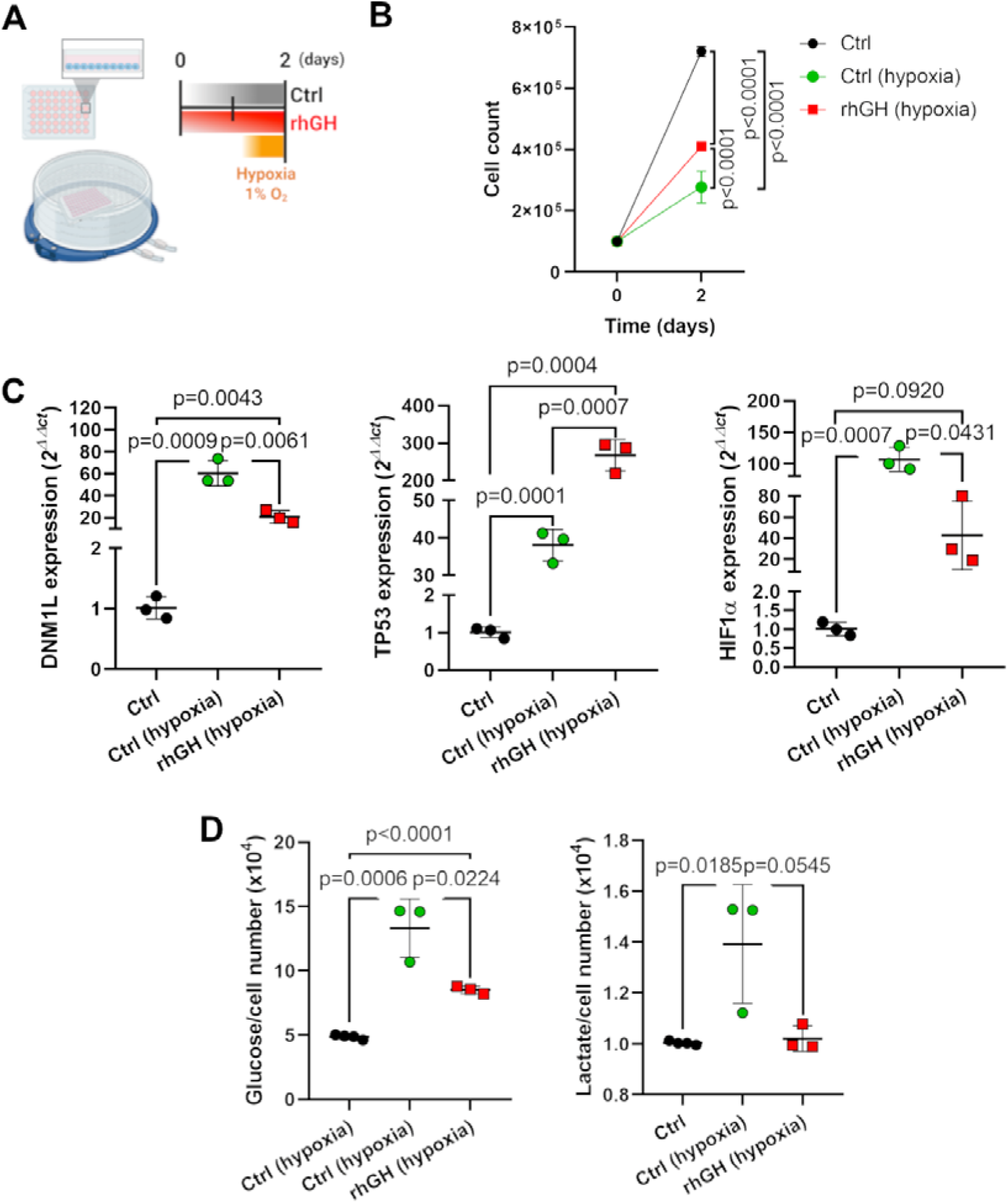
Growth hormone modulates proliferation, metabolic stress responses, and glucose metabolism under hypoxic conditions. (A) Schematic representation of experimental design. MDA-MB-231 cells were cultured under normoxic or hypoxic conditions (1% O□) for 2 days in the presence or absence of recombinant human growth hormone (rhGH). (B) Cell growth curves showing total cell number at day 0 and after 2 days of culture. (C) Relative mRNA expression of *DNM1L*, *TP53*, and *HIF1A* in control and hypoxia-exposed cells with or without rhGH. (D) Glucose consumption and lactate production measured in culture media after 2 days under hypoxia. Gene expression was quantified by qRT-PCR and normalized using the 2^−ΔΔCT^ method. Metabolite levels were normalized to cell number. Data are presented as mean ± SEM with individual points representing independent biological replicates. Statistical significance was determined using an unpaired Student’s *t*-test, with exact *p* values indicated.

### 6. Growth hormone accelerates tumor progression and metabolic remodeling in 3D breast cancer spheroids

To better mimic the solid tumor microenvironment and more realistically investigate the effects of growth hormone (GH) on tumor metabolism and progression, MDA-MB-231 mKate2 cells were cultured as three-dimensional (3D) spheroids using a bioprinting-based approach (Fig. 6A). The resulting spheroids displayed a uniform size distribution and a highly compact three-dimensional architecture, characterized by the presence of a hypoxic core enriched in non-viable (necrotic) cells and surrounded by a viable peripheral cell layer. In addition, spheroids exhibited stable and homogeneous red fluorescence throughout the structure, confirming sustained mKate2 expression (Fig. 6B-C). Quantitative analysis revealed that GH-treated spheroids exhibited increased total protein content (Fig. 6D), along with higher glucose consumption (Fig. 6E) after 4 days of exposure, indicating accelerated tumor progression, enhanced metabolic activity, and increased glycolytic flux. Notably, this effect contrasts with two-dimensional cultures, in which tumor progression becomes evident only after 6 days of GH treatment (Fig. 1B-C). In contrast, lactate production in GH-treated spheroids fluctuated over the exposure period and remained markedly low at day 4, differing from control spheroids, which showed a progressive increase in lactate production over time (Fig. 6F). Western blot and densitometric analyses revealed that GH treatment increased DRP1 phosphorylation (pDRP1) while maintaining MFN2 expression (Fig. 6G), suggesting enhanced mitochondrial remodeling dynamics. Notably, these effects mirror those observed in two-dimensional cultures, although they occur at later time points in the 3D spheroid context. In addition, acetyl-p53 levels were significantly reduced in GH-treated spheroids (Fig. 6G), consistent with the decreases observed in two-dimensional cultures at 2- and 4-day time points (Fig. 1G).

**Figure 6.**
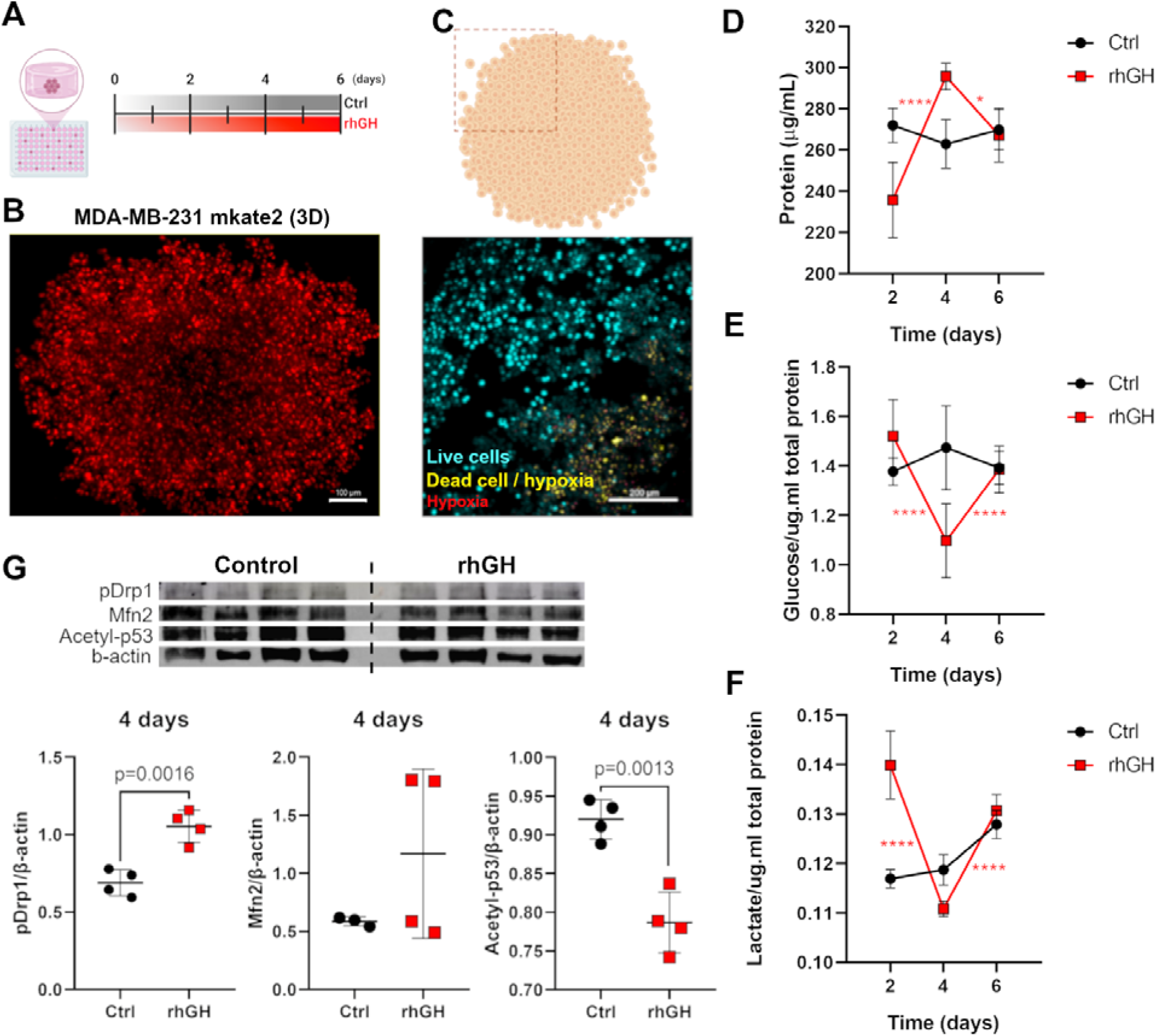
Growth hormone enhances metabolic activity and mitochondrial remodeling in 3D bioprinted breast cancer spheroids and promotes inflammatory signaling in zebrafish xenografts. (A) Experimental timeline showing the generation of 3D MDA-MB-231 spheroids and treatment with recombinant human growth hormone (rhGH) or vehicle control over a 6-day period. (B) Representative confocal image of 3D MDA-MB-231-mKate2 spheroids, illustrating tumor cell organization. (C) Representative fluorescence images of spheroid cross-sections showing live cells, dead cells, and hypoxic regions, highlighting spatial metabolic heterogeneity within 3D cultures. (D) Total protein content normalized to spheroid volume measured over time in control and rhGH-treated spheroids. (E) Glucose consumption normalized to total protein across the culture period. (F) Lactate production normalized to total protein over time. (G) Representative western blot analysis of mitochondrial dynamics and stress-response proteins, including phosphorylated DRP1 (pDRP1), MFN2, and acetyl-P53, with β-actin as a loading control. Data represent mean ± SEM; * *p*<0.05, \*\**p*<0.01, *** *p*<0.001, \*\*\*\**p*<0.0001. Statistical significance was determined using an unpaired Student’s *t*-test, with exact *p* values indicated.

### 7. Growth Hormone Enhances Tumor Engraftment and Promotes a Pro-Inflammatory Microenvironment

To evaluate whether GH-induced metabolic alterations influence tumor–host interactions, 2D- and 3D-derived GH-treated MDA-MB-231 cells were xenografted into adult zebrafish larvae expressing GH-GFP (Fig. 7A). Confocal imaging and western blot analyses confirmed successful tumor engraftment (Fig. 7B–D). GH exposure enhanced tumor cell signal intensity, an effect that was particularly pronounced in zebrafish implanted with 3D-derived spheroids. In addition to imaging-based evidence, we detected increased expression of human Lamin B1 within the tumor region of zebrafish exhibiting high GH expression that were implanted with 3D MDA-MB-231 cultures, further confirming successful injection and tumor establishment. Transcriptional analysis of zebrafish genes revealed that successfully xenografted animals displayed upregulation of pro-inflammatory genes, including *cxcr4b*, *il8*, and *il12* (Fig. 7E). Together, these findings indicate that GH-driven metabolic and mitochondrial remodeling in 3D breast cancer spheroids. GH exposure increased tumor cell signal intensity and elevated expression of human Lamin B1 within tumor regions, accompanied by upregulation of pro-inflammatory genes, including *cxcr4b*, *il8*, and *il12*.

**Figure 7.**
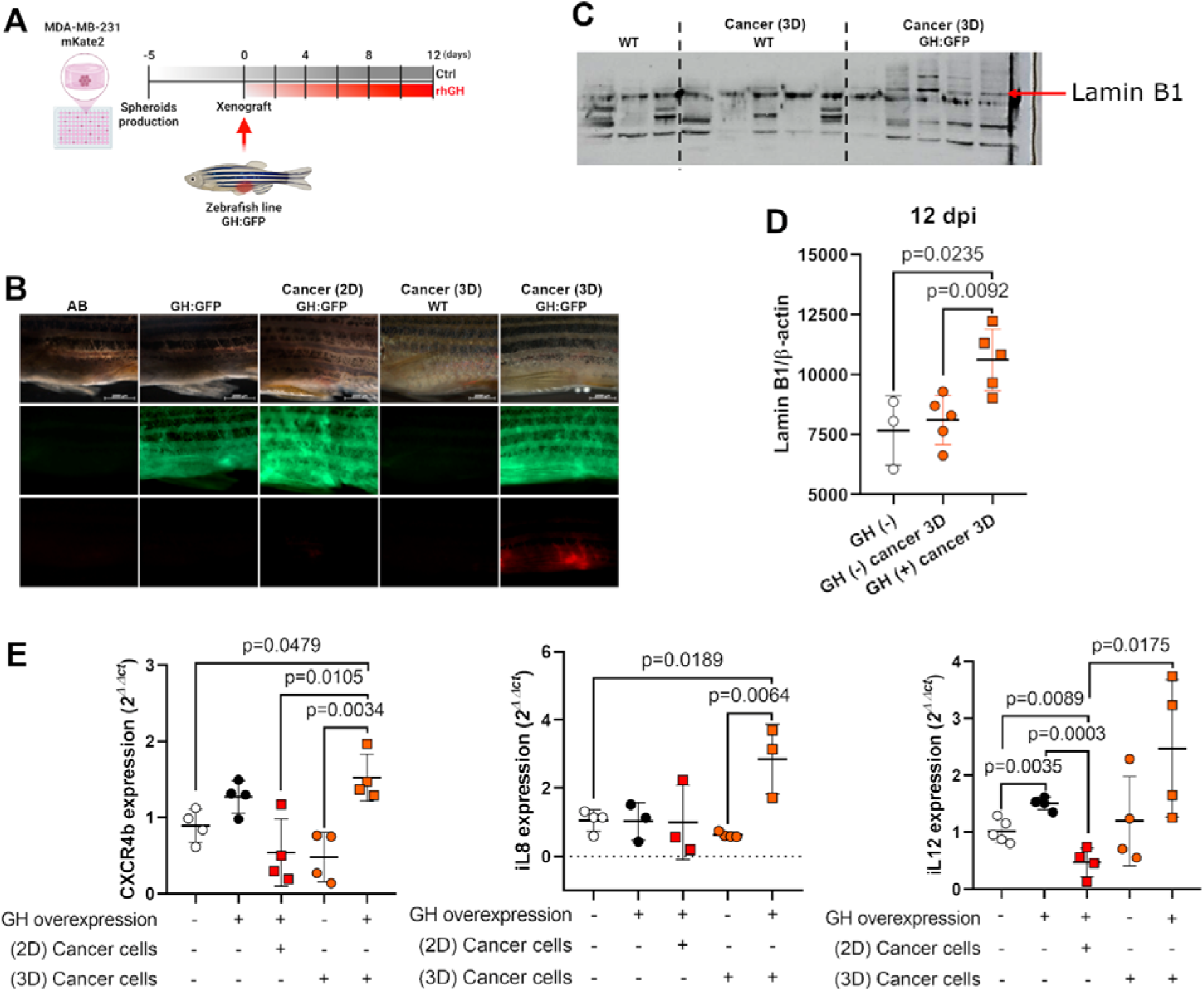
GH-conditioned MDA-MB-231 cells are associated with higher engraftment and increased host inflammatory gene expression. (A) Experimental schematic illustrating the production of 3D MDA-MB-231 spheroids, xenograft procedure, and timeline of recombinant human GH (rhGH) exposure in GH-GFP zebrafish. Spheroids were implanted at 0 days post injection (dpi), and animals were analyzed up to 12 dpi. (B) Representative brightfield and confocal images of zebrafish larvae showing tumor engraftment of 2D- and 3D-derived MDA-MB-231 cells in WT and GH-GFP hosts. Green fluorescence indicates host GH-GFP expression, and red signal corresponds to implanted tumor cells. (C) Representative western blot detecting human Lamin B1 in zebrafish tumor regions, confirming successful engraftment of human cancer cells. (D) Quantification of human Lamin B1 normalized to β-actin at 12 dpi, showing significantly increased. (E) Transcriptional analysis of zebrafish pro-inflammatory genes (*cxcr4b*, *il8*, and *il12*) by qPCR. Data represent mean ± SEM. Statistical significance was determined using an unpaired Student’s *t*-test or two-way ANOVA followed by Sidak’s post hoc test, with exact *p* values indicated.

### 8. Transcriptomic integration identifies conserved metabolic and inflammatory pathways associated with TNBC

To determine whether growth hormone–associated metabolic and stress-response signatures observed in our experimental models are reflected in human disease, we integrated publicly available transcriptomic datasets derived from breast cancer patients and TNBC cell models (Fig. 8A). Differential expression analyses were performed using the GSE36295 and GSE38959 patient cohorts, comparing normal breast tissue with TNBC samples, and the GSE250157 dataset derived from TNBC cell lines with differential HLA expressions. Intersection analysis revealed a shared set of genes commonly dysregulated across datasets, with 35 genes overlapping between all three cohorts (Fig. 8B), indicating a conserved transcriptional program associated with TNBC biology. Pathway enrichment analysis of the intersecting genes from the GSE36295 and GSE38959 datasets identified significant enrichment of cell-cycle–related pathways, including E2F targets and G2/M checkpoint signaling, as well as interferon-mediated inflammatory responses, mitotic spindle organization, and mTORC1 signaling (Fig. 8C). When intersecting all datasets, G2/M checkpoint, mitotic spindle and E2F tgargets pathways remained prominently enriched, suggesting robust conservation of proliferative programs across clinical samples and cellular models (Fig. 8D). Heatmap visualization of the GSE36295 cohort demonstrated clear segregation between normal and TNBC samples, with coordinated upregulation of genes related to mitochondrial dynamics, hypoxia, chemokine signaling, and stress responses, including *MFN2, TP53, HIF1A, CXCR4, CXCL8, CXCL12,* and *DNM1L* (Fig. 8E). Consistent with these observations, violin plot analysis confirmed significantly higher expression of *CXCR4* and *HIF1A* in TNBC samples compared with normal breast tissue (Fig. 8F). Finally, pathway enrichment analysis of the GSE38959 dataset revealed strong enrichment of MYC target gene sets, mitochondrial signaling, oxidative phosphorylation, glycolysis, epithelial–mesenchymal transition, interferon responses, and DNA repair pathways (Fig. 8G), highlighting the coexistence of metabolic reprogramming, inflammatory signaling, and proliferative stress in TNBC. Together, these results demonstrate that TNBC consistently shows cell-cycle and interferon activation, with cohort-dependent evidence for mitochondrial and metabolic pathway upregulation aligned with GH-driven phenotypes.

**Figure 8.**
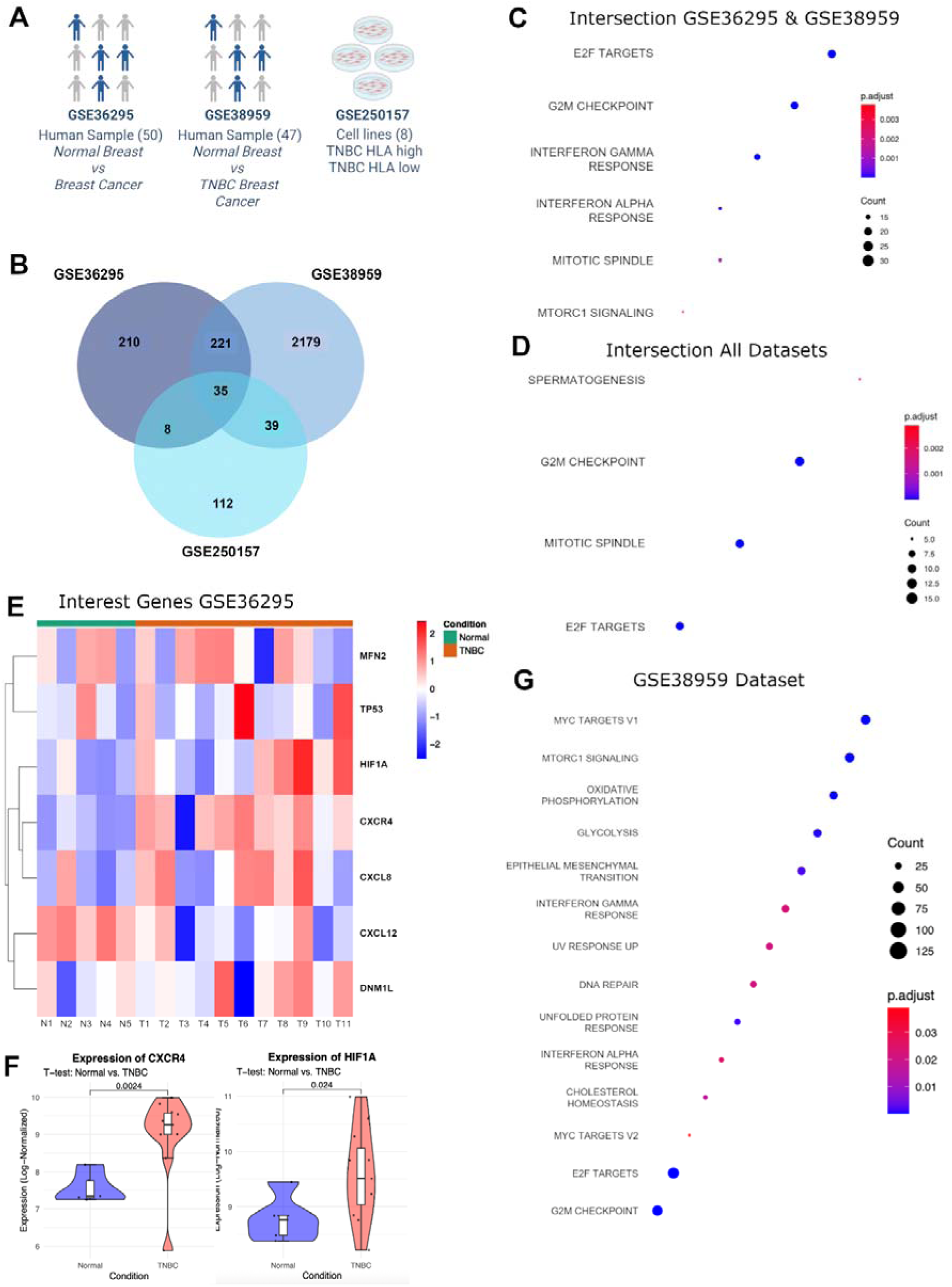
Integrated transcriptomic analysis identifies conserved proliferative and inflammatory pathways and cohort-dependent metabolic signals in TNBC. (A) Schematic overview of the transcriptomic datasets used in the analysis, including GSE36295 and GSE38959 human breast cancer cohorts comparing normal breast tissue and TNBC samples, and the GSE250157 TNBC cell line dataset stratified by HLA expression. (B) Venn diagram showing the overlap of differentially expressed genes among GSE36295, GSE38959, and GSE250157, identifying a shared core gene signature across patient samples and cellular models. (C) Pathway enrichment analysis of intersecting genes between GSE36295 and GSE38959, revealing significant enrichment of cell-cycle–related pathways (E2F targets, G2/M checkpoint), interferon signaling, mitotic spindle organization, and mTORC1 signaling. (D) Pathway enrichment analysis of genes shared across all three datasets, highlighting conserved activation of G2/M checkpoint and mitotic spindle pathways. (E) Heatmap of selected differentially expressed genes in the GSE36295 dataset, showing clear segregation between normal and TNBC samples and coordinated regulation of genes associated with mitochondrial dynamics, hypoxia, chemokine signaling, and stress responses. (F) Violin plots illustrating increased expression of CXCR4 and HIF1A in TNBC samples compared with normal breast tissue in the GSE36295 cohort. (G) Pathway enrichment analysis of the GSE38959 dataset demonstrating enrichment of MYC target genes, mitochondrial signaling, oxidative phosphorylation, glycolysis, epithelial–mesenchymal transition, interferon responses, and DNA repair pathways. Dot size represents gene count, and color indicates adjusted p value. Statistical significance thresholds are indicated in each analysis.

## Discussion

In this study, we show that growth hormone (GH) signaling engages DRP1-dependent mitochondrial remodeling to integrate metabolic plasticity and stress adaptation in triple-negative breast cancer (TNBC). GH increased pDRP1 and mitochondrial fragmentation, and both were reversed by Mdivi-1, supporting DRP1 involvement in the phenotype. Across complementary experimental systems, including 2D and 3D cultures, hypoxic conditions, in vivo xenografts, and integration with human transcriptomic datasets, GH engaged DRP1-mediated mitochondrial fission to coordinate glycolytic metabolism, proliferative capacity, and stress-response signaling. GH increased proliferation, protein content, and mitochondrial mass without increasing OCR under protein normalization, consistent with a regulated Warburg phenotype in which mitochondria support biosynthesis, redox homeostasis, and signaling rather than ATP production per se ^18–20^. These findings align with prior evidence that GH promotes TNBC growth and metastasis, whereas GHR inhibition suppresses proliferation and oncogenic signaling ^21,22^.

Although DRP1 overexpression is widely documented in cancer, its upstream regulation remains poorly defined. We show that GH promotes mitochondrial fission by increasing DRP1 abundance and phosphorylation, positioning DRP1 as a convergence point between GH signaling and mitochondrial architecture, consistent with its regulation by MAPK- and CDK-dependent growth factor pathways ^23^. Pharmacological inhibition of mitochondrial fission uncoupled GH-driven glycolytic activation from proliferative output, demonstrating a hierarchical relationship in which mitochondrial dynamics determine whether metabolic reprogramming is productively coupled to proliferation. These effects occurred without changes in respiratory capacity, reinforcing the concept that mitochondrial dynamics primarily regulate metabolic flexibility rather than oxidative output ^24^. DRP1-dependent mitochondrial remodeling also functions upstream of stress signaling. GH dynamically regulated TP53 and HIF1A, with early stress suppression followed by activation under sustained exposure, consistent with adaptive responses to metabolic pressure. Inhibition of mitochondrial fission by Mdivi-1 attenuated GH-induced TP53 and HIF1A upregulation and further reduced acetyl-p53 levels, indicating that mitochondrial dynamics modulate stress signaling through post-translational control of p53 activity ^15,25,26^. Given the central roles of TP53 and HIF1A as integrators of metabolic, hypoxic, and proliferative cues, these data position mitochondrial fission upstream of stress-response coordination.

Under hypoxia, GH retained its ability to enhance proliferation and glucose consumption, revealing a context-dependent metabolic rewiring in which lactate accumulation was stabilized rather than increased. This suggests that GH redirects glucose-derived carbons toward alternative metabolic or redox-supporting pathways to limit acidification under hypoxic stress, a defining feature of aggressive solid tumors ^14,27^. GH effects were amplified in 3D tumor spheroids, emerging earlier than in 2D cultures, underscoring the importance of tumor architecture in shaping metabolic and stress responses ^28^.

In vivo, GH-conditioned tumor cells, particularly those derived from three-dimensional cultures, exhibited enhanced engraftment and growth while inducing a pro-inflammatory host transcriptional program, including cxcr4b, il8, and il12, consistent with established roles of inflammatory chemokines in tumor–microenvironment interactions and niche formation ^29,30^. Given that mitochondrial dynamics and glycolytic metabolism regulate inflammatory signaling, ROS production, and chemokine expression, GH-driven mitochondrial remodeling is likely to contribute to the establishment of a tumor-permissive inflammatory niche ^6,31^.

Concordantly, integration of human TNBC transcriptomic datasets revealed enrichment of mitochondrial dynamics, glycolysis, hypoxia signaling, cell-cycle progression, and inflammatory pathways, with consistent upregulation of TP53, HIF1A, DNM1L, MFN2, and CXCR4, supporting a model in which GH-responsive mitochondrial–metabolic circuits are embedded within core TNBC biology. TP53 and HIF1A act as central stress-response hubs that integrate metabolic, hypoxic, and proliferative signals in cancer cells, coordinating adaptive transcriptional programs in response to environmental pressure ^14,32^. Under hypoxic or metabolic stress, stabilization of HIF1A promotes glycolytic reprogramming and pro-inflammatory, pro-migratory pathways, including IL8 and CXCR4 expression, thereby facilitating tumor–microenvironment interactions and invasive behavior ^16,29,33,34^. Findings derive from MDA-MB-231 (mutant TP53) and may not capture the full spectrum of TNBC heterogeneity; cross-line validation and isogenic analyses are warranted. Together, these findings support a pathway-centric model in MDA-MB-231 in which DRP1-dependent mitochondrial remodeling modulates GH activity, motivating validation across TNBC genotypes, acting as a critical downstream integrator that determines how GH-driven metabolic reprogramming is coupled to proliferation and stress adaptation.

While our data support a central role for the GH–DRP1 axis in regulating mitochondrial dynamics, glycolytic metabolism, and proliferation in TNBC, several limitations should be acknowledged. Inferences regarding DRP1 dependence are primarily based on pharmacologic inhibition using Mdivi-1; orthogonal approaches, including genetic loss-of-function and rescue experiments, will be required to establish definitive causality. We also did not directly quantify cell-cycle entry under DRP1 inhibition; future work incorporating DNA-content profiling and S-phase and mitotic markers will be important to assess coupling between metabolic reprogramming and cell-cycle progression. Finally, given reports of potential off-target effects of Mdivi-1, independent genetic perturbations of DNM1L and/or alternative DRP1 inhibitors will be required to establish direct causality, including for the modulation of TP53- and HIF1A-associated pathways.

## Conclusion

In conclusion, our study identifies DRP1-dependent mitochondrial remodeling as a critical downstream effector of growth hormone (GH) signaling that links metabolic reprogramming to proliferation and stress adaptation in triple-negative breast cancer. These data implicate mitochondrial dynamics as potential modulators of glycolytic control, stress signaling, and tumor–microenvironment interactions, these findings reveal a previously unrecognized mechanism through which endocrine cues shape tumor aggressiveness. Together, this work highlights mitochondrial fission as a central determinant of GH-driven metabolic plasticity and suggests that targeting the GH–DRP1 axis may represent a strategy to limit tumor adaptability and progression in TNBC.

## Acknowledgments

This research was supported by the São Paulo Research Foundation (FAPESP; grants 2017/05264-7, 2021/03192-4, 2023/07482-2), the National Council for Scientific and Technological Development, and the Coordenação de Aperfeiçoamento de Pessoal de Nível Superior–Brasil (finance code 001).

## Authors’ Contributions

J.M.M.G. conceived and designed the study, performed experiments, analyzed and interpreted the data, and wrote the manuscript.

M.T.P. and L.M.S. performed experiments, collected data, and contributed to data analysis.

L.E.D.G. performed the bioinformatic analyses and contributed to data interpretation.

M.A.A., L.C.P., C.M.F., and B.N.P. assisted with experimental procedures and data collection.

M.C.C. contributed to experimental design and provided technical support.

N.O.S.C. supervised the study, provided resources and infrastructure, and contributed to the interpretation of the results and manuscript revision.

All authors reviewed, edited, and approved the final version of the manuscript.

## Ethics approval and consent to participate

All animal experiments were approved by the Ethics Committee for Animal Use (CEUA-ICB/USP; protocol no. 9710200720) and conducted in accordance with the Brazilian Guidelines for Care and Use of Animals for Scientific and Teaching Purposes (CONCEA) as well as international guidelines for animal experimentation. Publicly available datasets were used in this study; therefore, no ethical approval or informed consent was required.

## Consent for publication

Not applicable.

## Conflict of interest statement

The authors declare no conflicts of interest.

## Notes

### Competing Interest Statement

The authors have declared no competing interest.

