## Supplementary figures and images for "DRP1-mediated mitochondrial fission integrates growth hormone signaling with metabolic and stress adaptation in triple-negative breast cancer"

### Figure S1

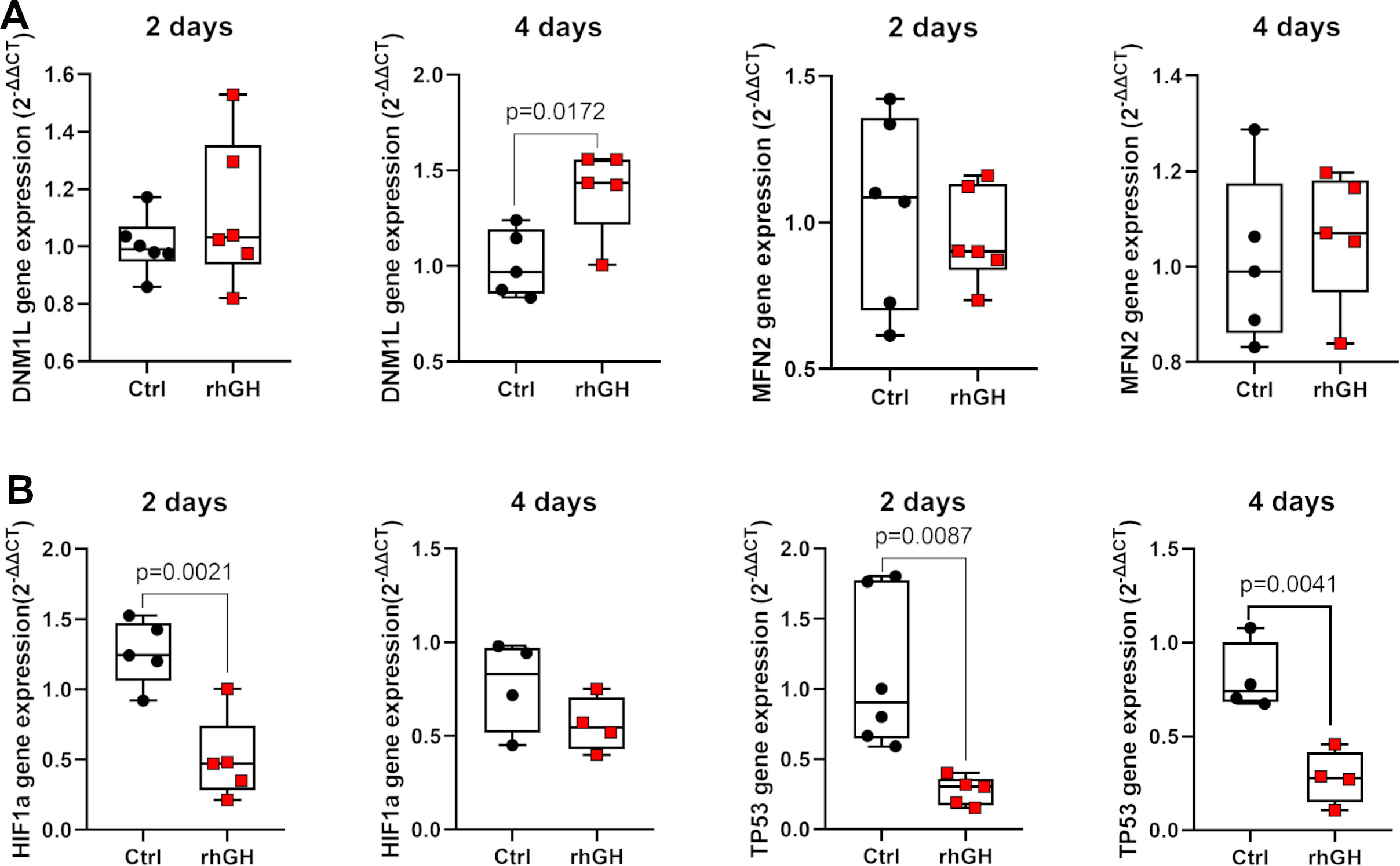

### Figure S2

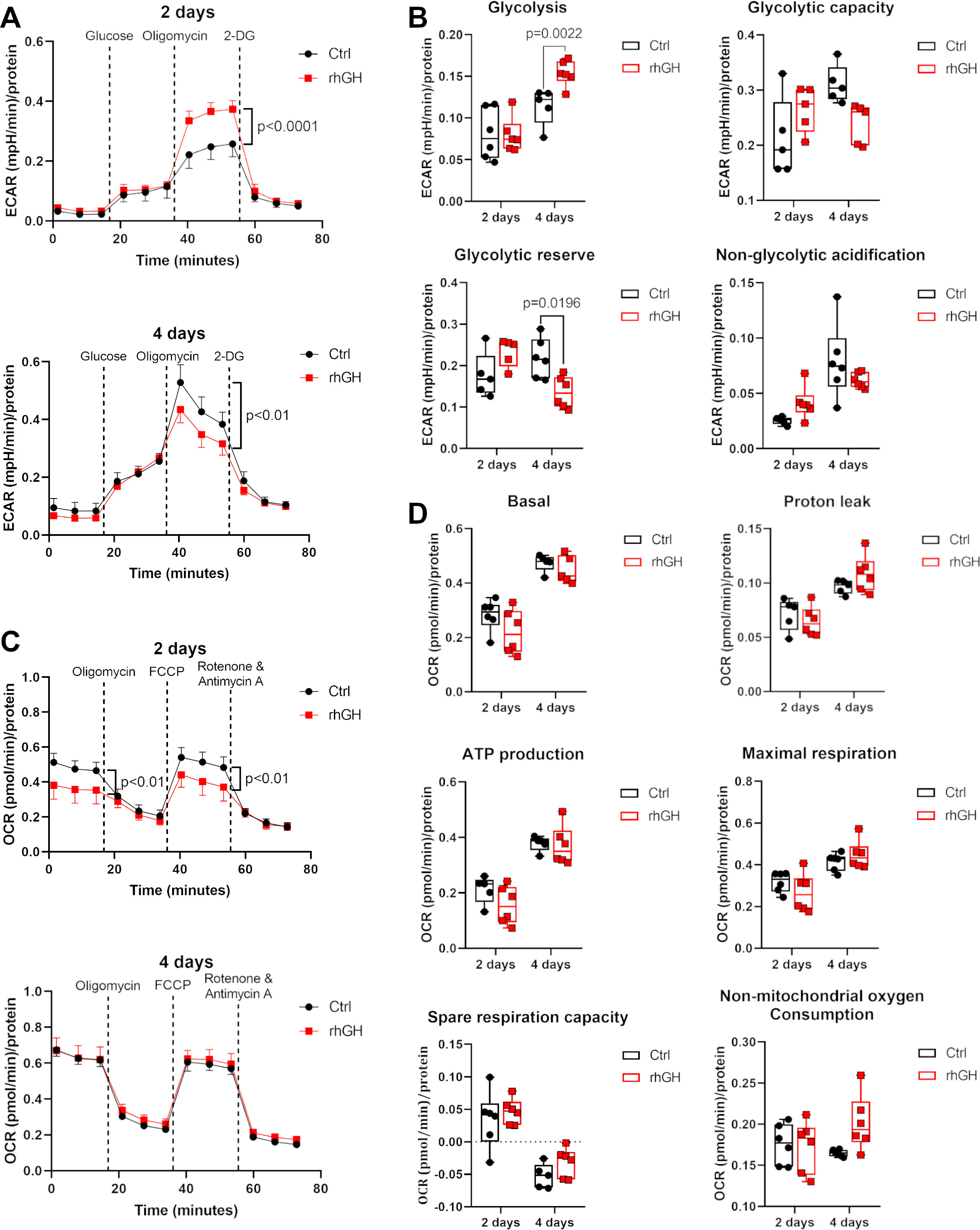

### Figure S3

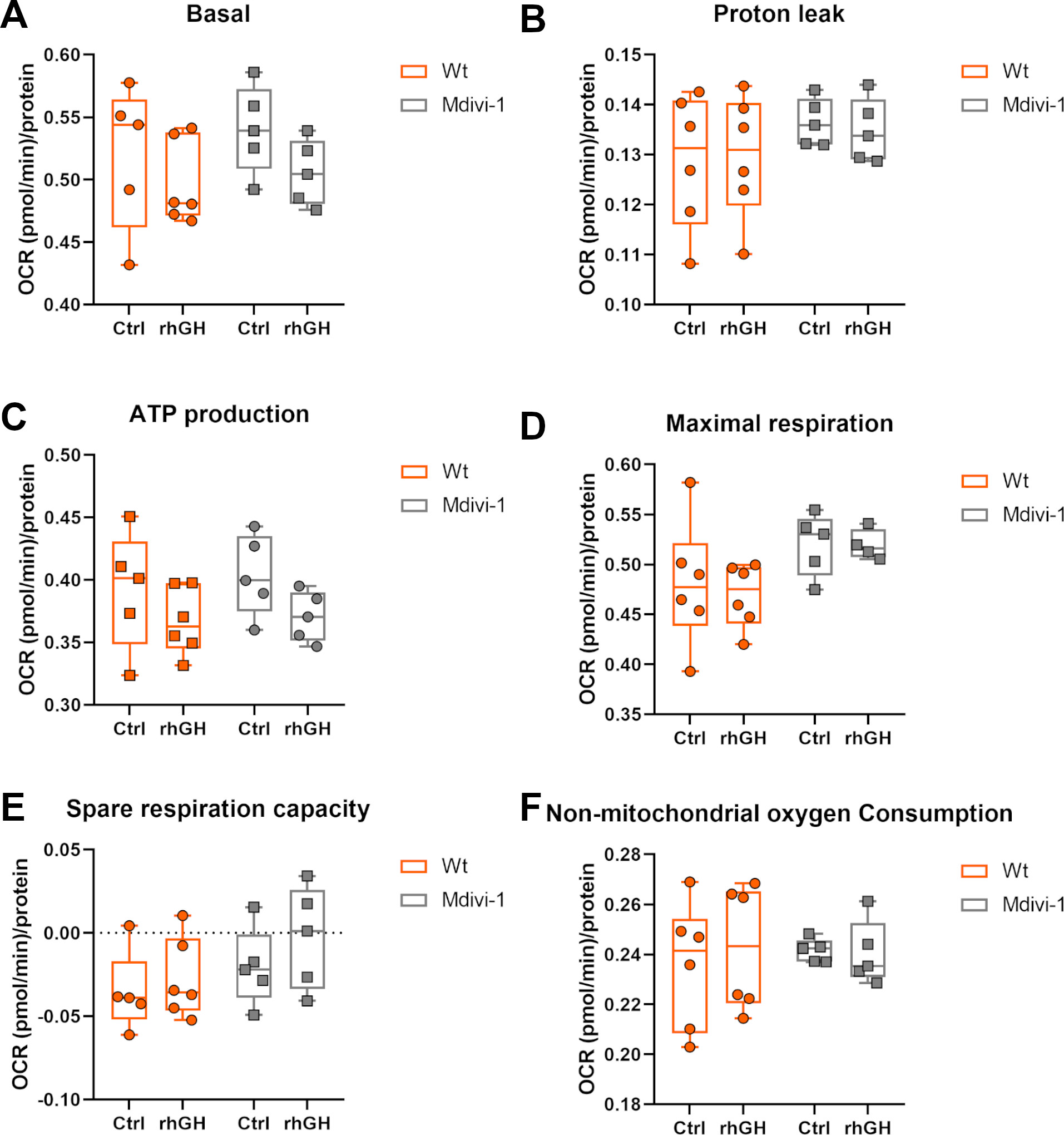

### Figure S4

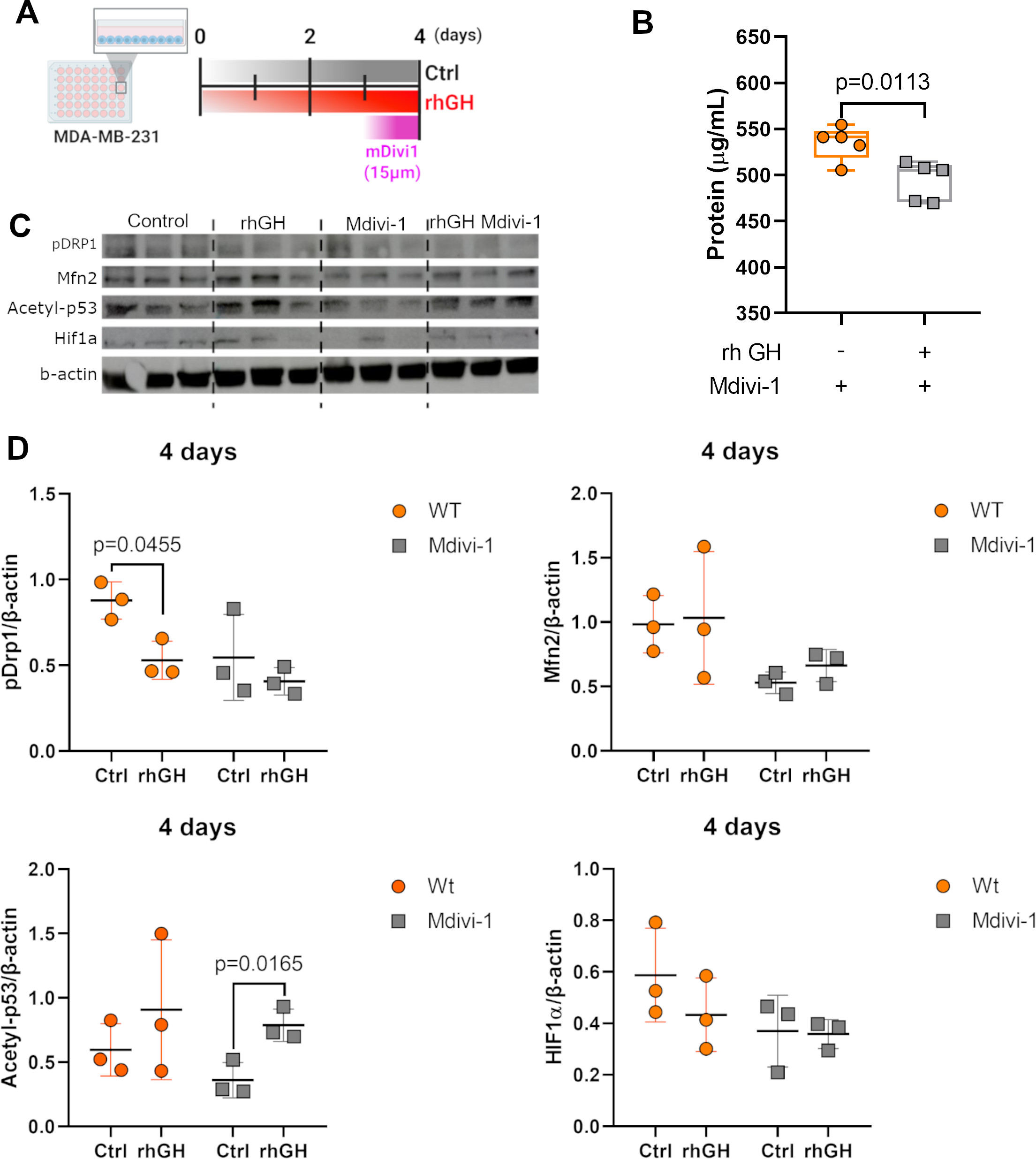
