## Supplementary material for "DRP1-mediated mitochondrial fission integrates growth hormone signaling with metabolic and stress adaptation in triple-negative breast cancer": Table 1

### **Supplementary Table 1. Primer sequences used for qPCR analysis**

| Gene | Direction | Sequence (5'–3') |
| --- | --- | --- |
| Human DNM1L | Forward | GATGCCATAGTTGAAGTGGTGAC |
| Human DNM1L | Reverse | CCACAAGCATCAGCAAAGTCTGG |
| Human MFN1 | Forward | GGTGAATGAGCGGCTTTCCAAG |
| Human MFN1 | Reverse | TCCTCCACCAAGAAATGCAGGC |
| Human HIF1A | Forward | GAAAGCGCAAGTCTTCAAAG |
| Human HIF1A | Reverse | TGGGTAGGAGATGGAGATGC |
| Human ACTB | Forward | CATGTACGTTGCTATCCAGGC |
| Human ACTB | Reverse | CTCCTTAATGTCACGCACGAT |
| Human TP53 | Forward | ATGGAGGAGCCGCAGTCAGAT |
| Human TP53 | Reverse | ACCTGGGTCTTCAGTGAACCATTG |
| Human RPLP0 | Forward | AGCCCAGAACACTGGTCTC |
| Human RPLP0 | Reverse | ACTCAGGATTTCAATGGTGCC |
| Zebrafish il12a | Forward | ACGCAAACGGTGTCTGTCT |
| Zebrafish il12a | Reverse | CTCTGTAGGCATTCGCTCTCAT |
| Zebrafish il8 | Forward | GTTTTCCTGGCATTTCTGACCA |
| Zebrafish il8 | Reverse | GCGTCGGCTTTCTGTTTCAA |
| Zebrafish cxcr4b | Forward | CTTACTTACCCGCAGGAGGC |
| Zebrafish cxcr4b | Reverse | CCAGGCAGCAAAAAGCCAAT |
