## Supplementary Figure Legends for "DRP1-mediated mitochondrial fission integrates growth hormone signaling with metabolic and stress adaptation in triple-negative breast cancer"

### **Supplementary Figure S1.** Growth hormone modulates the expression of mitochondrial dynamics and stress-response genes in breast cancer cells. (A) Relative mRNA expression of *DNM1L* and MFN2, genes involved in mitochondrial fission and fusion, respectively, after 2 and 4 days of culture in control conditions (Ctrl) or in the presence of recombinant human growth hormone (rhGH). (B) Relative mRNA expression of stress-response genes *HIF1A* and *TP53* after 2 and 4 days of culture with or without rhGH. Gene expression was quantified by qRT-PCR and normalized to housekeeping genes using the 2^−ΔΔCT^ method. Data are shown as box-and-whisker plots with individual data points representing independent biological replicates. Statistical significance was determined using an unpaired Student’s t-test; p values are indicated in the graphs.

**Supplementary Figure S2.** Time-dependent effects of growth hormone on glycolytic and mitochondrial function in breast cancer cells. (A) Representative extracellular acidification rate (ECAR) profiles of control (Ctrl) and recombinant human growth hormone–treated (rhGH) MDA-MB-231 cells after 2 and 4 days of culture. (B) Quantification of glycolytic parameters derived from ECAR measurements, including basal glycolysis, glycolytic capacity, glycolytic reserve, and non-glycolytic acidification at 2 and 4 days. (C) Representative oxygen consumption rate (OCR) profiles of Ctrl and rhGH-treated cells at 2 and 4 days. (D) Quantification of mitochondrial respiration parameters derived from OCR measurements, including basal respiration, proton leak, ATP-linked respiration, maximal respiration, spare respiratory capacity, and non-mitochondrial oxygen consumption at 2 and 4 days. ECAR and OCR values were normalized to total protein content. Data are presented as box-and-whisker plots with individual data points representing independent biological replicates. Statistical significance was determined using a two-way ANOVA followed by Sidak’s post hoc test, and an unpaired Student’s t-test; *p* values are indicated in the graphs.

**Supplementary Figure S3.** DRP1 inhibition modulates mitochondrial respiration in growth hormone–treated breast cancer cells. (A–F) Quantification of mitochondrial respiration parameters derived from oxygen consumption rate (OCR) measurements in wild-type (Wt) MDA-MB-231 cells and cells treated with the DRP1 inhibitor Mdivi-1, cultured in control conditions (Ctrl) or in the presence of recombinant human growth hormone (rhGH). Parameters shown include (A) basal respiration, (B) proton leak, (C) ATP-linked respiration, (D) maximal respiration, (E) spare respiratory capacity, and (F) non-mitochondrial oxygen consumption. OCR values were normalized to total protein content. Data are presented as box-and-whisker plots with individual points representing independent biological replicates. Statistical significance was determined using two-way ANOVA followed by Sidak’s post hoc test, with exact *p* values indicated.

### **Supplementary Figure S4.** Early effects of DRP1 inhibition during GH exposure. (A) Experimental timeline illustrating MDA-MB-231 cell culture and treatment conditions. Cells were cultured for four days in the presence or absence of recombinant human growth hormone (rhGH). The DRP1 inhibitor Mdivi-1 (15 µM) was added during the final 24 hours following three days of GH exposure. (B) Total protein content (µg/mL) after 4 days of treatment, as an indicator of cell proliferation following GH and Mdivi-1 treatments. (C) Representative immunoblots showing phosphorylated DRP1 (pDRP1), MFN2, acetyl-P53, and HIF-1α protein levels under the indicated conditions. β-actin was used as a loading control. (D) Quantification of protein levels normalized to β-actin after four days of treatment. Data are shown as individual values with mean ± SEM. Statistical significance was determined using two-way ANOVA followed by Sidak’s post hoc test, and an unpaired Student’s t-test with exact *p* values indicated.
